# Checkpoint proteins in patients with precancer and cervical cancer

**DOI:** 10.1101/2021.02.09.430409

**Authors:** Elena Kayukova, Leonid Sholokhov, Tatiana Belokrinitskaya, Pavel Tereshkov

**Author notes:** ^#^Corresponding author. These authors contributed equally to this work. The efficiency of immuno-oncological drugs in treatment cervical cancer is not high. The research of new immunological targets in cervical cancer is actual. **The aim** was to study the local level of checkpoint proteins of immune cycle in patients with cervical cancer of different stages (sCD25, 4-1BB, B7.2, TGF-b1, CTLA-4, PD-L1, PD-1, Tim-3, LAG-3, Galectin-9, sCD27, PD-L2). **Materials and research methods.** The objects of study were the patients with precancer (n=13) and cervical cancer (n=49). The control group consisted of 13 gynecologically healthy women. The material of study was cervical epithelium obtained by cervical brush. The research method was flow cytometry. Test parameters: sCD25, 4-1BB, B7.2, TGF-b1, CTLA-4, PD-L1, PD-1, Tim-3, LAG-3, Galectin-9, sCD27, PD-L2. Statistical analysis of the data was carried out using nonparametric statistics using the Mann – Whitney U-test. **The results of the study.** The levels of sCD25, B7.2, PDL1, Tim-3, sCD27, PD-L2 were increased in the comparison groups relation to the control (p <0.05). The differential criterion for invasive cervical cancer may be an increase in the level of LAG-3 in the cervical epithelium by 43% (p <0.05).The levels of PD-L1, PD-1, sCD27 and PD-L2 were increased opposite the levels LAG-3 and Galectin-9 were decreased in patients with metastatic cervical cancer(p <0.05).

## Abstract

The efficiency of immuno-oncological drugs in treatment cervical cancer is not high. The research of new immunological targets in cervical cancer is actual.

The aim was to study the local level of checkpoint proteins of immune cycle in patients with cervical cancer of different stages (sCD25, 4-1BB, B7.2, TGF-b1, CTLA-4, PD-L1, PD-1, Tim-3, LAG-3, Galectin-9, sCD27, PD-L2).

Materials and research methods. The objects of study were the patients with precancer (n=13) and cervical cancer (n=49). The control group consisted of 13 gynecologically healthy women. The material of study was cervical epithelium obtained by cervical brush. The research method was flow cytometry. Test parameters: sCD25, 4-1BB, B7.2, TGF-b1, CTLA-4, PD-L1, PD-1, Tim-3, LAG-3, Galectin-9, sCD27, PD-L2. Statistical analysis of the data was carried out using nonparametric statistics using the Mann – Whitney U-test. The results of the study.

The levels of sCD25, B7.2, PDL1, Tim-3, sCD27, PD-L2 were increased in the comparison groups relation to the control (p <0.05). The differential criterion for invasive cervical cancer may be an increase in the level of LAG-3 in the cervical epithelium by 43% (p <0.05).The levels of PD-L1, PD-1, sCD27 and PD-L2 were increased opposite the levels LAG-3 and Galectin-9 were decreased in patients with metastatic cervical cancer(p <0.05).

## INTRODUCTION

The efficiency of immunotherapy of patients with metastatic cervical cancer (MCC) is not high [1], known difficulties of objective assessment of PD-L1 expression as a predictor of tumor response to immunotherapy [2] are background causes for continuous search for pathogenetically substantiated additional immunological targets of cervical cancer (CC) cells and tumor microenvironment.

## EXPERIMENTAL PART

**Aim of the research**: to assess levels of certain immune checkpoint proteins in CC patients (sCD25, 4-1BB, B7.2, TGF-b1, CTLA-4, PD-L1, PD-1, Tim-3, LAG-3, Galectin-9, sCD27, PD-L2) according to the spread of the neoplastic process.

### Materials and methods

We performed a non-randomized controlled prospective study among patients with precanceros conditions and CC. The studied groups similar in age and associated diseases (average age was 39±9.8 years old) were:

I clinical group - patients with precancerous cervical diseases - cervical intraepithelial neoplasia III degree (n=13).
II group - initial non-treated patients with squamous CC of I-IV stages (n=49), of whom patients with non-invasive (NI) n=8, invasive local (IL) n=18, local spread (n=19), metastatic CC n=4.

The control group included 13 gynecologically healthy volunteers, who were informed with the design of the research, and who gave informed consent to participate in it (average age was 30.0±4.4). The research was carried out in compliance with the principles of the WMA Declaration of Helsinki, 1964, 2013 R.E), and upon approval of the Local committee on ethics of the Chita State Medical Academy, Russian Federation (protocol 44 dated 09.11.2012).

The material of study was cervical epithelium, collected by standard method by taking with cytobrush. The method of flow cytometry FC500 (Beckman Coulter, USA) with the use of reagents included in the panel HU Immune Checkpoint Panel 1 - S/P (10-plex) w/FP was used to identify parametrs: sCD25, 4-1BB, B7.2, TGF-b1, CTLA-4, PD-L1, PD-1, Tim-3, LAG-3, Galectin-9, sCD27, PD-L2.

Statistical processing of data was computer-based with application of BIOSTAT software by nonparametric statistics methods using the Mann – Whitney U-test. The differences were considered statistically significant at p<0.05.

## RESULTS

The results of intergroup analysis are given in Table 1.

**Table 1.**
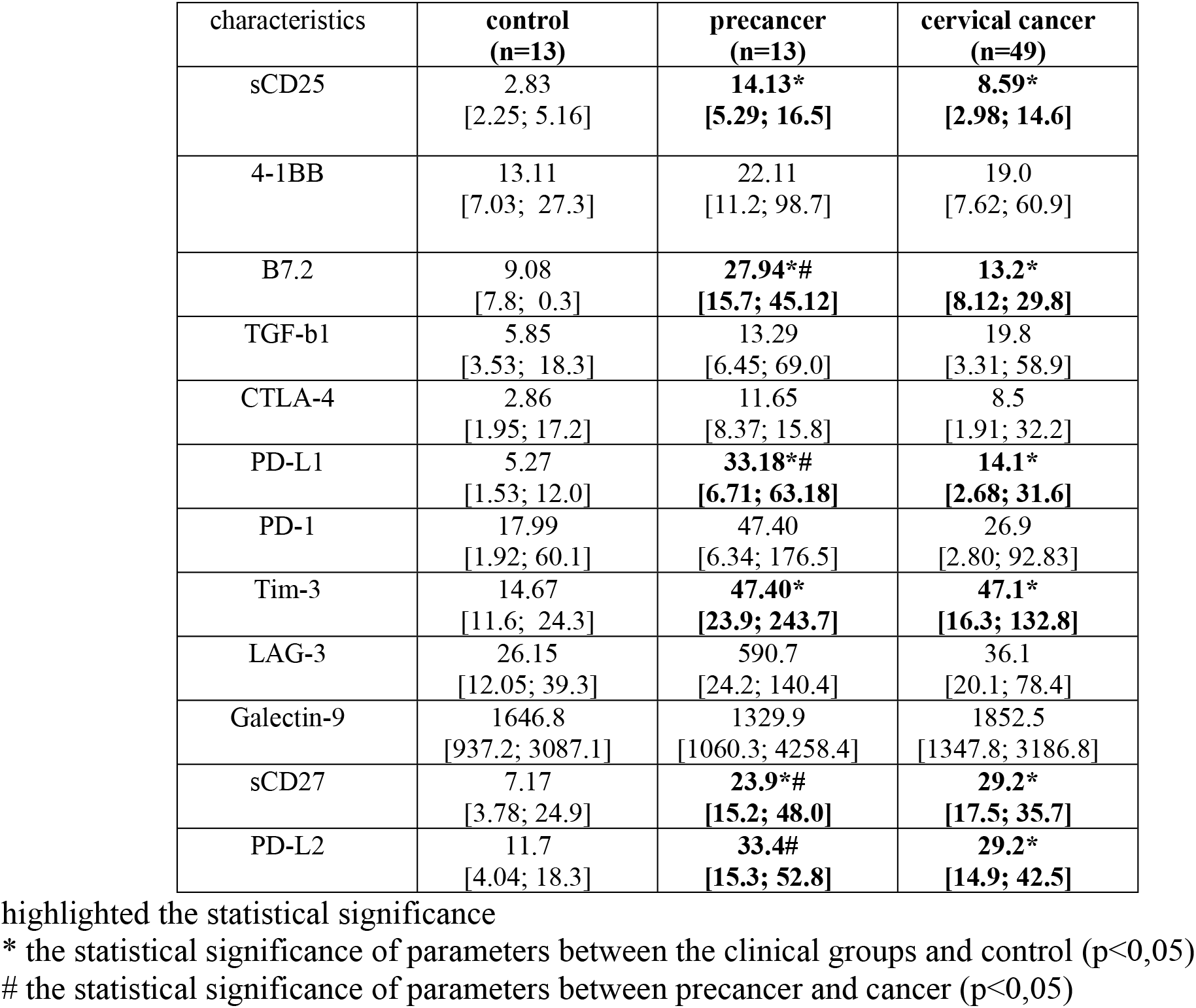
The local level of immune checkpoint proteins in patients with the damage of uterine cervix Me (Q1; Q3), pg/ml

| characteristics | **control**<br>**(n=13)** | **precancer**<br>**(n=13)** | **cervical cancer**<br>**(n=49)** |
| --- | --- | --- | --- |
| sCD25 | 2.83<br>[2.25; 5.16] | **14.13\***<br>**[5.29; 16.5]** | **8.59\***<br>**[2.98; 14.6]** |
| 4-1BB | 13.11<br>[7.03; 27.3] | 22.11<br>[11.2; 98.7] | 19.0<br>[7.62; 60.9] |
| B7.2 | 9.08<br>[7.8; 0.3] | **27.94\*#**<br>**[15.7; 45.12]** | **13.2\***<br>**[8.12; 29.8]** |
| TGF-b1 | 5.85<br>[3.53; 18.3] | 13.29<br>[6.45; 69.0] | 19.8<br>[3.31; 58.9] |
| CTLA-4 | 2.86<br>[1.95; 17.2] | 11.65<br>[8.37; 15.8] | 8.5<br>[1.91; 32.2] |
| PD-L1 | 5.27<br>[1.53; 12.0] | **33.18\*#**<br>**[6.71; 63.18]** | **14.1\***<br>**[2.68; 31.6]** |
| PD-1 | 17.99<br>[1.92; 60.1] | 47.40<br>[6.34; 176.5] | 26.9<br>[2.80; 92.83] |
| Tim-3 | 14.67<br>[11.6; 24.3] | **47.40\***<br>**[23.9; 243.7]** | **47.1\***<br>**[16.3; 132.8]** |
| LAG-3 | 26.15<br>[12.05; 39.3] | 590.7<br>[24.2; 140.4] | 36.1<br>[20.1; 78.4] |
| Galectin-9 | 1646.8<br>[937.2; 3087.1] | 1329.9<br>[1060.3; 4258.4] | 1852.5<br>[1347.8; 3186.8] |
| sCD27 | 7.17<br>[3.78; 24.9] | **23.9\*#**<br>**[15.2; 48.0]** | **29.2\***<br>**[17.5; 35.7]** |
| PD-L2 | 11.7<br>[4.04; 18.3] | **33.4#**<br>**[15.3; 52.8]** | **29.2\***<br>**[14.9; 42.5]** |

highlighted the statistical significance
\* the statistical significance of parameters between the clinical groups and control ( $p<0,05$ )

In the patients of the I clinical group the levels of sCD25, B7.2, PDL1, Tim-3, sCD27 were 5; 3; 6,3; 3,2 and 3,3 times higher than the corresponding control values (p<0,05). In the II clinical group an increase of control values of sCD25, B7.2, PDL1, Tim-3, sCD27, PD-L2 by 3; 1,5; 2,7; 3,2; 4 and 2.5 times accordingly was identified (p<0,05). During intergroup analysis increased level of indicators B7.2, PD-L1, PD-L2 (by 2,1; 2,4; 1,1 times accordingly (p<0,05)) with decreased sCD27 by 1.2 times (p<0,05) was identified in the I group.

The results of subgroup analysis of CC patients according to the spread of the neoplastic process are given in Table 2.

**Table 2.** Level of immune checkpoint proteins in cervical mucus of CC patients in accordance with the spread of the neoplastic process Me (Q1; Q3), pg/ml

| characteristics | control group<br>(n=13) | noninvasive cancer<br>(n=8) | invasive local<br>cancer<br>(n=18) | local spread cancer<br>(n=19) | metastatic cancer<br>(n=4) |
| --- | --- | --- | --- | --- | --- |
| sCD25 | 2.83<br>[2.25; 5.16] | 6.85<br>[2.76; 13.3] | **11.2\***<br>**[4.57; 15.3]** | **13.8\***<br>**[3.0; 12.2]** | 9.8<br>[2.29; 9.8] |
| 4-1BB | 13.11<br>[7.03; 27.3] | 27.3<br>[15.1; 46.1] | **56.0\***<br>**[11.3; 81.6]** | **15.5#**<br>**[7.57; 19.9]** | 169.9<br>[103.4; 266.8] |
| B7.2 | 9.08<br>[7.8; 0.3] | 13.6<br>[8.57; 16.1] | 22.6<br>[8.28; 55.0] | 16.4<br>[8.08; 24.1] | 23.5<br>[18.1; 33.7] |
| TGF-β1 | 5.85<br>[3.53; 18.3] | 14.7<br>[3.44; 34.5] | 44.0<br>[12.5; 64.8] | **18.6#**<br>**[3.20; 27.1]** | 33.1<br>[28.9; 58.8] |
| CTLA-4 | 2.86<br>[1.95; 17.2] | [13.65<br>[3.53; 36.3] | 17.8<br>[5.52; 37.7] | **21.4#**<br>**[1.79; 18.4]** | **2.27\***<br>**[2.27; 2.93]** |
| PD-L1 | 5.27<br>[1.53; 12.0] | 15.0<br>[3.44; 28.9] | **22.5\***<br>**[5.25; 37.2]** | **25.6#**<br>**[2.37; 23.4]** | **48.6^**<br>**[26.8; 72.1]** |
| PD-1 | 17.99<br>[1.92; 60.1] | 58.3<br>[19.8; 105.5] | 73.0<br>[9.87; 109.1] | **34.7#**<br>**[1.85; 28.2]** | **193.9^**<br>**[142.0; 332.0]** |
| Tim-3 | 14.7<br>[11.6; 24.3] | 75.6<br>[19.8; 140.1] | **74.6\***<br>**[37.6; 147.8]** | **59.2#**<br>**[14.4; 59.2]** | **52.8\***<br>**[44.6; 65.5]** |
| LAG-3 | 26.15<br>[12.05; 39.3] | **34.3<sup>v</sup>**<br>**[21.1; 58.6]** | 58.9<br>[27.5; 95.7] | **541.2#**<br>**[11.3; 42.8]** | **377.3^**<br>**[377.3; 739.5]** |
| Galectin-9 | 1646.8<br>[937.2; 3087.1] | 1724.0<br>[1347.8; 1818.8] | 2003.0<br>[1153.7; 3148.6] | 1997.1<br>[1428.6; 3673.9] | **1313.7^**<br>**[1129.8; 1413.9]** |
| sCD27 | 7.17<br>[3.78; 24.9] | **18.6\***<br>**[11.9; 24.9]** | **20.4\***<br>**[17.2; 26.5]** | **31.6\*#**<br>**[17.5; 52.5]** | **97.6\*^**<br>**[66.8; 128.4]** |
| PD-L2 | 11.7<br>[4.04; 18.3] | 15.9<br>[11.3; 36.0] | **28.9\***<br>**[19.8; 36.6]** | **29.2\***<br>**[17.5; 57.5]** | **66.9\*^**<br>**[58.5; 75.4]** |

highlighted the statistical significance ( $p<0,05$ )
\* the statistical significance of parameters between the clinical groups and control
<sup>v</sup> - the statistical significance of parameters between noninvasive cancer and invasive local cancer

When we analyzed the data depending on the spread of the neoplastic process we did not identify any significant differences between the levels of B72, TGF-b1, PD-1, Galectin-9, LAG-3 as compared to the control values (p>0,05).

The level of sCD25 was increased in patients with invasive local and local spreadCC by 4 and 5 times accordingly as compared to the control value (p< 0.05).

Changes in the values of 4-1BB and PD-L1 were identified only in the subgroup of patients with invasive local CC: the increase was by 4.3 times as compared with the control value for each of them (p< 0,05).

It was established that only the patients with metastatic CC had 20% decreased level of CTTLA4 (p< 0,05).

The value of Tim-3 in the cervical smear of patients with invasive local and metastatic CC increased the control value by 5 and 3.6 times accordingly (p< 0,05).

The levels of sCD27 и PD-L2 increased in all subgroups of CC patients with the maximum levels being observed in patients with metastatic CC.

Subgroup analysis of indicators of patients with invasive local and local spreadCC showed decreased level of 4-1BB, TGF-b1, PD-1 and Tim-3 (by 3.77 (p<0.05), 2.4 (p<0.05); 2.1 (p<0.05) и 1.3 (p<0.05) times accordingly) with increased levels of CTLA4, PD-L1, LAG-3, (by 1.2 (p<0.05); 1.1 (p<0.05), 9 (p<0.001); 1.5 and 1 times (p<0.001) accordingly).

In the group of patients with metastatic CC as compared with local spreadCC significant differences of PD-L1, PD-1, LAG-3 and Galectin-9 were identified, whereas the value of the first two was increased by 1.9 (p<0.05) and 5.6 (p<0.05) times accordingly and the value of the last ones was decreased by 1.4 (p<0.05) and 1.5 times (p<0.05) accordingly.

## DISCUSSION

**sCD25** is one of the regulators of immune reactions in autoimmune, oncological diseases and critical conditions of intensive care patients [3, 4]. sCD25 is produced under the influence of a number of ferments (elastase, MMP-9) during shedding, which leads to inactivation of immune cells, in particular to functional deficiency of NK-cells, and to suppression of anti-tumor activity of T-cells which intensifies tumor progression [3]. Pathogenetic mechanisms of sCD25 interaction with immune cells are being studied. It should be noted that we have not found any data on the study of sCD25 levels in CC patients. However, our data conforms with the results of other research of other neoplasia [5]. High serum level of sCD25 is an indicator of worse survival rates and unresponsiveness to immunotherapy of patients with melanoma. Sh. Abdelfattah et al. suggested that identification of sCD25 in blood serum should be used as a marker of hepatocellular carcinoma taking into account the detected relationship between its quantity and disease stage.

**Receptor 4-1BB** is a member of TNF receptor family, expressed on CD8, CD4- T-cells, B-lymphocytes, regulatory T-lymphocytes, NK cells, dendritic cells, mast cells, etc. Its physiological role is to regulate activity of the above mentioned cells by activating a cascade of signaling events (JNK, ERK, β-catenin and AKT) [6]. 4-1BB promotes survival of T-cells, participates in the formation of immunological memory by activating antiapoptopic genes Bcl-2, Bcl-xl, и Bfl-1, induces proliferation of T-cells and enhances their effector function. Activation of 4-1BB leads to aging of dendritic cells, enhances their survival and production of cytokines IL-6, IL-12 и IL-27 [7]. Dual capacity of 4-1BB to stimulate reaction of effector T-cells on pathogenes with simultaneous limitation of autoimmune reactions made this receptor an alluring target for tumor immunotherapy. There were published results of research studying 4-1BB agonists in the therapy of some tumors [8]. T. Bartkowiak et al. showed in vitro efficiency of administration of intranasal peptide vaccine HPV E6/E7, which contained αGalCer as an adjuvant in combination with systemic administration of agonistic antibodies to treat HPV-driven tumor [9]. We have not found any literature showing data on the study of 4-1BB levels in CC patients. Our data on the maximum level of 4-1BB in patients with local spreadCC suggests tense activation of one of the elements of antitumor immune response.

**Ligand B7.2** is expressed on antigen presenting cells (APCs) and can interact with T-lymphocytes receptor CD28, inducing their activation and triggering immune response, or with activated T-lymphocytes receptor CTLA4 blocking immune reactions. In the absence of costimulation of CD28 T-cells affected by the antigen become anergic. Due to insufficient expression B7-2 cells are unable to transmit co-stimulatory signals required for efficient activation of T-cells, which potentiates tumor progression [10]. Pathogenetic participation of B7 family proteins in carcinogenesis has been proven for pulmonary cancer, hemato-oncological diseases, stomach cancer, ovarian cancer, clear-cell carcinoma [11]. We are the first to study local level of B7.2 in patients with cervical disease. High level of B7.2 in both comparison groups is a reflection of activation of T-cell antitumor immune response. It’s interesting to note the discovered fact of decreased level of B7.2 in CC patients by 2.1 times (p <0,05) as compared to the corresponding indicators of patients with precancerous cervical conditions, which can indicate reduced efficiency of immune reactions during cervical carcinogenesis.

**TGF-b1** (Transforming growth factor beta) acts as one of the key regulators of the development and progression of tumor. It influences all important stages of cancer progression including migration and invasion. Paradoxical effect of TGF-β in the neoplastic process has been established. At the initial stages of tumor development, it gives antineoplastic action, but during its progression its effect becomes co-carcinogenic. This TGF-β effect is explained by changing activity of micro-RNA which control TGF-β activity, TGF-β gene polymorphism, condition of endogenous proteins [12]. Pathogenetic role of TGF-β was established in carcinogenesis of colorectal cancer, urogenital cancers, breast cancer. In our research we have not received any significant differences in the value of TGF-β among the studied groups [12].

**CTLA-4** (cytotoxic T-lymphocyte-associated protein 4) is a negative immune regulator which suppresses both antiviral and antitumor immune response. CTLA4 integrates with co-stimulatory molecules CD80 (B7-1) and CD86 (B7-2) expressed on the surface of antigen presenting cells (APCs) and suppress T-lymphocyte function. The data on the research of CTLA4 expression in CC patients is scarce. Meta-analysis of 11 international studies among 3899 cases of CC showed significant connection between the development of tumor in carriers of polymorphism rs5742909 gene CTLA4 among Asians [13]. Abnormal expression of CTLA-4 in T-lymphocytes isolated from peripheral blood of metastatic CC patients is known, which is related to genetic defects in the gene that codes a corresponding protein, as well as with a low level of its antagonist with immune stimulating action - CD28, which is associated with reciprocal interaction [14]. Increased expression of CTLA-4 on circulating T-lymphocytes reflects exhaustion of T-cells due to continuous antigen stimulation, and simultaneous identification of CD28 expression in CD8+ subset of T-cells is a marker that this process can be reversed [15]. CTLA-4 can as well be expressed on the surface of T-regulatory cells and enhance its immunosuppressive action in case of distributed CC [16]. The data we received on the extremely low CTLA4 level in metastatic CC patients coincide with the conclusions of researchers headed by A. Gutiérrez-Hoya (2019), who established low expression of CTLA4 by cells of experimental CC-Hela cell line [17].

**PD-1** (Programmed cell death-1 protein) is a transmembrane protein which is expressed by T-lymphocytes. One of its ligands is **PD-L1**, which is expressed on the surface of tumor cells of cervix, APCs, and T-lymphocytes. During interaction between PD-1 and PD-L1 suppression of antitumor and antiviral immune response is observed. Information on PD-1 and PD-L1 expression is numerous and contradictory. Generally, 46-61% of initial CC express PD-1 and 22-96% - PD-L1 [18]. There is an assumption on the influence of HPV contamination on the expression of PD-L1 and PD-1 [19]. Analysis of PD-1 and CTLA4 expression in 33 various morphological types of tumor has been published. PD-1 and CTLA4 expression varies in different types of tumor and is not characteristic for squamous CC. High level of PD-1 expression in tumor samples of CC patients correlated with the best disease-free survival (p = 0.047) [20]. Such conclusions were received by R Karim [21]. R. Grochot and al. reported the absence of prognostic levels of PD-L1 and PD-1 expression in CC patients [22]. Y. Meng et al. published information on detected correlation between PD-L1 expression in CC patients with clinical signs of disease progression and unfavorable morphological prognosis factors: metastatic lesions of lymph nodes and lymph vessel invasion [23]. Y. Liu et al reported unfavorable prognosis role of high levels of PD-L1 and PD-1 in patients with recurrent and metastatic CC which corresponds to worse disease free and general survival of patients [18].

In order to give prognosis on the course of neoplastic process not only the level of immune response inhibitors expression is important, but also the number of immune cells located in the locus of cancer and at the periphery. All this can be assessed by a prognostic immune scoring system called Immunoscore. H. Chen et al. were the first to apply Immunoscore to CC patients and established that a favorable prognostic factor for CC patients was a positive PD-L1 expression on tumor cells surrounded by a large number of tumor-infiltrating T-lymphocytes [24].

Our data on the local levels of PD-L1 and PD-1 among the studied groups conform with the results of research by L. Mezache et al [19]. PD-L1 expression on tumor cells was less in case of squamous CC (51%) as compared to CIN 1-2 (95%). However, the level of PD-L1 expression on T-lymphocytes was the maximum in CC patients (80% versus 61%). The maximum values of PD-L1 and PD-1 in metastatic CC patients that we discovered is a mechanism for inhibiting antitumor immune response.

**PD-L2** is a second ligand for PD-1 receptor. It is expressed on APCs, and fibroblasts. PD-L2 as well as PD-L1 expression in normal tissues is limited. However, during carcinogenesis activation of PD-L2 / PD-1 path leads to the suppression of antitumor immune response which intensifies cancer growth [25]. Participation of PD-L2 / PD-1 in carcinogenesis is scarcely studied [27]. The role of PD-L2 is known in intensifying cancer progression (by activating Rhoa-ROCK-LIMK2, PI3K / AKT / mTOR paths), formation of tumor chemoresistance. Information on possible prognostic role of PD-L2 expression levels for some oncological diseases has been published [26].

**Tim-3** (T-cell immunoglobulin-3) is a protein, which is expressed on T-lymphocytes, and tumor cells, interacts with ligand – Galectin 9 and inhibits antitumor immune response. Tim-3 is seen as a negative regulator molecule important for the creation of immune tolerance in tumor microenvironment. Tim-3 and ligand PD-1 expression in T-cells leads to exhaustion of T-cell element and tumor growth [27]. Tim-3 activates IL-6-STAT3 path, which promotes metastasis of various tumors. The results of Y. Cao et al. conform with our data on the high level of Tim-3 in precancerous and cancerous lesions of cervix. The authors established correlative relationship between high level of Tim-3 and the level of neoplastic process, worse general survival, and metastatic potential of tumor [28]. The research of L. Zhang et al. established epigenetic regulation of Tim-3 expression in CC under the influence of E6 and E7 HPV proteins [29, 30].

**LAG-3** (lymphocyte-activation gene 3) is another protein which belongs to immune response inhibitors. LAG-3 is expressed on cell membranes TIL, activated CD4+ и CD8+ T-cells, and regulatory T-cells, NK, B-lymphocytes, dendritic cells. LAG-3 is connected with the main major histocompatibility complex-II on APCs, thus downregulating antitumor immune response. In the literature we found a reference to only one research which indicated LAG-3 overexpression in HPV-infected CC cells, which conforms with our data [31].

**Galectin 9** (Gal-9) belongs to the family of beta-galactoside-binding proteins. Its pathophysiological role in the carcinogenesis is complex. It is known that Gal-9 exhibits proapoptotic effect by activating NK and p38-MAPK paths, Caspase 3, −8, − 9, which was demonstrated on myeloma cells. Gal-9 inhibits invasion and metastasis of myeloma tumor cells and hepatocarcinoma by influencing the epithelial–mesenchymal transition. The receptor for Gal-9 on immune cells is Tim-3. Their interaction leads to reduction of anti-tumor activity of immune cells [33]. Sporadic publications report low expression of Gal-9 on CC tumor cells [33]. With increase in the level of dysplastic changes in the cervical epithelial cells the level of Gal-9 expression decreases, which conforms to the significantly low level of Gal-9 in metastatic CC patients, that we established.

**sCD27** is a soluble form of transmembrane protein CD27 (T-lymphocyte receptor), that serves as a co-stimulator of immune reactions. Its ligand is CD70 expressed on immune cells. Their interaction promotes stimulation of immune response. Pathogenetic role of sCD27 has been established for the development of autoimmune (systemic lupus erythematosus), viral (HIV infection) and some oncological diseases (Hodgkin lymphoma, prostate cancer, lung cancer, breast cancer, colon cancer, glioblastoma, etc.) [34, 35, 36]. We have not found any literature showing data on the study of sCD27 in CC patients. The information about increased amount of sCD27 in the cervical mucus of patients with precancerous and oncological cervical diseases indicate activation of cell-bound immunity. However, when we cross-check this data with clinical information, we can conclude that sCD27 activation is not sufficient to inhibit progression of CC. It may be connected with blockage of sCD27 ligand expressed not on T-lymphocytes but on different cells which interferes with the activation of antitumor immune response.

## CONCLUSIONS

1. It was found that patients with precancerous diseases and CC patients have increased levels of sCD25, B7.2, PDL1, Tim-3, sCD27, PD-L2 in the cervical mucus as compared with control value (p<0,05). Significant increase in the levels of B7.2, PD-L1 and PD-L2 was found in the patients with precancerous diseases by 2.1; 2,2, and 1.1 times respectively relative to the comparison groups (p<0,05).
2. Differential criterion of invasive CC can be 43% increased level of LAG-3 in cervical mucus (p<0,05) compared to the corresponding indicator in patients with non-invasive CC.
3. Metastatic CC by contrast with local spreadCC is characterized by increased levels of PD-L1, PD-1, sCD27 and PD-L2 by 1.9 (p<0.05); 5.6 (p<0.05); 3 (p<0.001) and 2.3 times (p<0.001), correspondingly with decreased values of LAG-3 and Galectin-9 by 1.4 (p<0.05) and 1.5 times accordingly (p<0.05).
4. Invasive local CC by contrast with local spreadCC is characterized by decreased levels of 4-1BB, TGF-b1, PD-1 and Tim-3 (by 3.77 (p<0.05), 2.4 (p<0.05); 2.1 (p<0.05) и 1.3 (p<0.05) times accordingly) with increased levels of CTLA4, PD-L1, LAG-3, sCD27 (by 1.2 (p<0.05); 1.1 (p<0.05), 9 times (p<0.001); 1.6 times accordingly (p<0.001).

## CONCLUSION

In the course of cervical carcinogenesis expression of immune proteins that have a key role in the balanced immune cell responses is interrupted on the local level. It is possible that the identified changes have a prognostic character in relation to the progression of the disease, and have a predictive value for an efficient immunotherapy of CC patients, which requires further research. The authors declare that there is no conflict of interests.

The research was carried out within the state research assignment for 2018-2020 (№AAAA-A17-117030310232-5).

